# Dopamine enhances GABA_A_ receptor-mediated current amplitude in a subset of intrinsically photosensitive retinal ganglion cells

**DOI:** 10.1101/2023.12.11.571141

**Authors:** Nikolas Bergum, Casey-Tyler Berezin, Jozsef Vigh

**Author notes:** Correspondence: Jozsef Vigh.

## Abstract

Neuromodulation in the retina is crucial for effective processing of retinal signal at different levels of illuminance. Intrinsically photosensitive retinal ganglion cells (ipRGCs), the neurons that drive non-image forming visual functions, express a variety of neuromodulatory receptors that tune intrinsic excitability as well as synaptic inputs. Past research has examined actions of neuromodulators on light responsiveness of ipRGCs, but less is known about how neuromodulation affects synaptic currents in ipRGCs. To better understand how neuromodulators affect synaptic processing in ipRGC, we examine actions of opioid and dopamine agonists have on inhibitory synaptic currents in ipRGCs. Although µ-opioid receptor (MOR) activation had no effect on γ-aminobutyric acid (GABA) currents, dopamine (via the D1R) amplified GABAergic currents in a subset of ipRGCs. Furthermore, this D1R-mediated facilitation of the GABA conductance in ipRGCs was mediated by a cAMP/PKA-dependent mechanism. Taken together, these findings reinforce the idea that dopamine’s modulatory role in retinal adaptation affects both non-image forming as well as image forming visual functions.

## INTRODUCTION

Neuromodulators tune neuronal excitability throughout the central nervous system to regulate mood, reward, movement and cognition (1). In the retina, there is significant evidence that neuromodulators are crucial for both image-forming and non-image forming visual functions (2–9). Indeed, neuromodulators do alter the excitability and light responsiveness of melanopsin-expressing intrinsically photosensitive retinal ganglion cells (ipRGCs) (2, 7), the principal source of light information to areas of the brain that control non-image forming visual functions (10). Perhaps most significantly, the M1 subtype of ipRGCs is thought to transmit signals to the brain’s sleep/circadian centers, which is critical for the synchronization of circadian rhythms with environmental light/dark cycles (10). Thus, to better understand how light signal is transmitted from the environment to the brain’s sleep/circadian circuitry, it is important to develop a mechanistic understanding for how light-driven ipRGC activity might be altered by local neuromodulatory inputs. Moreover, an improved understanding of how neuromodulators regulate ipRGC function could provide valuable information that could help explain certain disorders and pathologies associated with altered ipRGC signaling, including the drug-mediated disruption of sleep/wake cycles.

Recent studies have highlighted the role that neuromodulatory inputs play in tuning ipRGC-mediated light-driven processes such as circadian photoentrainment and pupillary light reflex (2–12). Dopamine, the best studied neuromodulator in the retina, is considered a chemical messenger for light adaptation (9), by participating in the transition from rod-to cone-mediated retinal circuits and setting the gain of retinal neurons for optimal signaling in bright light (4, 13). The sole source of retinal dopamine are dopaminergic amacrine cells (DACs), which tonically release dopamine within the retina (4, 9, 13). Dopamine release increases during daylight; in contrast, a subset of cholinergic amacrine cells release endogenous opioid peptide β-endorphin during darkness (11, 14). Interestingly, it has been suggested that opioid receptors agonist (such as β-endorphin) suppress dopamine release (15) by activating µ-opioid receptors (MORs) expressed by DACs (16). Importantly, ipRGCs are known to express both MORs (2) and D1-type dopamine receptors (D1Rs) (7).

Previously our lab has shown that µ-opioid receptor (MOR) activation attenuated ipRGC light responses and decreased the excitability of ipRGCs (2). Interestingly, dopamine has also been shown to diminish light responses in these ganglion cell photoreceptors via D1Rs (7, 8). Taken together, direct neuromodulatory inputs to ipRGCs have a significant impact on ipRGC function. Studies examining the direct effect of dopamine and opioid agonists on ipRGC activity have been performed in dissociated ipRGC preparations or in the presence of synaptic blockers (2, 7). However light-evoked signaling of ipRGCs stems from the interaction of an intrinsic light response and rod and cone photoreceptor-driven visual signals that impinge on ipRGCs via excitatory and inhibitory synapses (17–19). Both opioid and dopamine signaling have been shown to alter synaptic communication in the nervous system, often via acting on the presynaptic terminals within the mesolimbic dopamine system (20–22). Nonetheless, whether synaptic inputs to ipRGCs are subject to opioid and/or dopaminergic modulation is not known. Thus, to develop a more complete understanding for how opioids and dopamine influence ipRGC activity, we examined how MOR and dopamine receptor activation alter synaptic inputs to ipRGCs.

Although very little is known about fast synaptic inputs to ipRGCs, results from the present study confirm earlier research reporting that only a subset of ipRGCs in the whole mount retina have detectable spontaneous synaptic events (18, 19). While spontaneous events were relatively small and infrequent, puffed γ-aminobutyric acid (GABA), but not glutamate, produced robust currents in enzymatically-dissociated solitary ipRGCs. D1R activation enhanced GABA currents in a subset of ipRGCs, whereas MOR activation had no effect on the amplitude of GABA puff-evoked currents. Our present results show that D1Rs facilitate GABAergic inputs to ipRGCs via a cAMP/PKA-dependent mechanism.

## MATERIALS AND METHODS

### Animals

We used transgenic Tg(*Opn4-*EGFP)ND100Gsat/Mmucd strain, generated by the GENSAT project. This strain carries a bacterial artificial chromosome (BAC) in which the melanopsin (*Opn4*) promoter drives expression of enhanced green fluorescent protein (EGFP); these mice are referred to as *Opn4*::EGFP (2, 23). Our study employed adult male and female animals for the experiments between the ages of 2-8 months. *Opn4*::EGFP mice were anesthetized with isoflurane and euthanized via cervical dislocation. Adult male and female animals (8+ weeks) were housed under a 12-h light:12-h dark cycle. They were fed standard chow and water ad libitum. All animals used in these studies were handled in compliance with the Institutional Animal Care and Use Committees of Colorado State University (Protocol 18-8395A, 28 January 2019) and in accordance with the ARVO Statement for the Use of Animals in Ophthalmic and Vision Research.

### Dissociated ipRGC preparation for whole cell recording

Solitary ipRGCs were enzymatically dissociated from retinas harvested from *Opn4*::EGFP+ mice as previously described (2, 24, 25). In brief, eyes from *Opn4*::EGFP+ mice were enucleated and hemisected posterior to the limbus; the lens and vitreous humor were removed. Retinas were detached from the retinal pigmented epithelium and incubated for 15 min at 37°C in a solution containing papain (10 U/ml, Worthington; Lakewood, NJ) and L-Cysteine hydrochloride (∼1 mg/mL, Sigma-Aldrich; Saint Louis, MO). After rinsing 3+ times in papain-free culture media, manual trituration was performed with a large-bore fire-polished Pasteur pipette and dissociated cells were plated on poly-D-lysine/laminin coated coverslips (Corning™BioCoat™; Bedford, MA) followed by >2 hours incubation in MACS® NeuroMedium without L-Glutamine (Miltenyi Biotech; Auburn, CA). The medium was supplemented with MACS® NeuroBrew-21, antibiotics (100 u/ml penicillin and 100 μg/ml streptomycin) and Gibco™ GlutaMAX™ Supplement (ThermoFischer; Waltham, MA, USA). Coverslips were maintained for 1-4 days in an incubator at 37°C before ipRGCs were targeted for patch clamp recordings.

### Whole mount retina preparation for whole cell recording

Whole mount retinas were harvested from *Opn4*::EGFP+ mice as described above (2). Following mouse euthanasia and enucleations, retinas were detached from the retinal pigment epithelium and placed in bicarbonate-buffered Ames’ medium (US Biological; Salem, MA) gassed with 95% O_2_/5% CO_2_ at room temperature. Retina sections were secured with a tissue anchor (harp) to the glass bottom of a superfusion chamber with the ganglion cell layer up. Recordings were made from these whole mount preparations within the same day of harvest.

### Whole cell patch clamp electrophysiology

M1 ipRGCs were identified and targeted based on their larger size (∼10 µm) and bright green fluorescence in either dissociated or whole mount retina preps made from *Opn4*::EGFP+ mice and studied in a perfusion chamber mounted on an upright microscope (Akioskop 2 FS plus, Zeiss; Oberkochen, Germany). Retina preparations were superfused at 2–5 ml/min with 300 ± 10 mOsmol bicarbonate-buffered Ames’ medium (US Biological; Swampscott, MA) constantly gassed with 95% O2 / 5% CO2 and were viewed through a 40x water immersion objective, with infrared differential contrast, and an infrared camera (AxioCam MRm, Zeiss; Oberkochen, Germany) with 2.5 pre-magnification ( Optovar, Zeiss; Oberkochen, Germany), which directed output to a 19” monitor (Westinghouse; Santa Fe Springs, CA).

A model p-97 horizontal puller (Sutter; Novato, CA) was used to pull patch pipettes from 1.5-mm-diameter, thick-walled borosilicate glass (World Precision Instruments; Sarasota, FL) which were then fire polished to a final resistance of 7–20 MΩ using a MF-830 Microforge (Narishige; Tokyo, Japan). Whole-cell voltage-clamp recordings were made from EGFP+ ipRGC somas using an EPC-10 USB patch-clamp amplifier and Patchmaster software version 2.3 (HEKA; Lambrecht (Pfalz), Germany) at room temperature during the light phase of the light/dark cycle. Membrane current and voltage data were filtered at 3 kHz. Recordings with leak >50 pA at −70 mV holding potential and/or series resistance (Rs) >50 MΩ at any time during the recording were terminated and excluded from analysis. Similarly, if the Rs changed more than 10% during the recording, there data were not considered for further analysis. The holding current to set the holding potential at −70 mV at break in was determined in voltage-clamp mode.

For GABA puff recordings, a cesium gluconate (CsGlc)-based internal solution was used that contained (in mM) the following: 100 Cs-gluconate, 10 phosphocreatine-di(tris) salt, 10 L-ascorbic acid, 2 EGTA, 3 Mg-ATP, 0.5 Na-GTP, 10 tetraethylammonium chloride, 0.1 CaCl2, 10 NaCl, pH 7.2 (adjusted with CsOH) and osmolarity of 295 ± 5 mOsmol.

To record spontaneous inhibitory events in the whole mount retina, a CsGlc-based internal solution was used that contained (in mM) the following: 110 Cs-gluconate, 10 phosphocreatine-di(tris) salt, 10 L-ascorbic acid, 2 EGTA, 3 Mg-ATP, 0.5 Na-GTP, 3 tetraethylammonium chloride, 2 QX314, 5 NaCl, pH 7.3 (adjusted with CsOH) and osmolarity of 295 ± 5 mOsmol. The standard extracellular solution was a bicarbonate-buffered Ames’ medium (US Biological; Salem, MA), with an osmolarity of 300 ± 10 mOsmol, continuously gassed with 95% O_2_ / 5% CO_2_. To record spontaneous inhibitory events, cells were held at +0 mV and recorded for 2-3 minutes. All recordings were performed in light adapted tissue.

### Immunohistochemistry

Immunohistochemistry was performed as previously described (11, 26, 27). Mice were anesthetized with isoflurane (Fluriso, VetOne) and sacrificed by cervical dislocation. The eyes were removed and dissected in 0.1 M phosphate buffered saline (PBS; pH 7.4). The cornea and lens were removed, and the resulting eyecups were fixed in freshly prepared 4% paraformaldehyde (PFA) in PBS then immersed in 30% sucrose at 4°C for several days prior to being embedded in Optimal Cutting Temperature (O.C.T.) Compound (Tissue-Tek). The tissue was sectioned at 20 μm on a ThermoFisher CryoStar NX50 cryostat, directly mounted on SuperFrost Plus slides (Fisher Scientific), and stored at −20°C until use. For each mouse, 3 or 4 slides, containing a total of 9 or 12 sections, respectively, were processed. The tissue was incubated in a blocking solution (5% normal goat serum and 0.5% Triton X-100 in PBS) for 1 h at RT, then in the anti-dopamine receptor antibody diluted in the blocking solution for 5-6 days at 4°C (rabbit anti-dopamine D1A receptor, 1:100, Sigma-Aldrich AB1765P). Sections were washed in PBS and incubated for 2 h at RT in the secondary antibody (goat anti-rabbit Alexa Fluor 555). Retinas were stained with DAPI (GeneTex GTX16206) and mounted in Vectashield Plus Antifade Mounting Medium (Vector Laboratories; Burlingame, CA).

All images were taken with a LSM 900 confocal microscope (Carl Zeiss; Oberkochen, Germany) with a 20× air objective. ipRGCs in the peripheral retina were identified based on positive GFP signal. Z-stack images were taken at 1 μm increments throughout the thickness of the sections. All images were ∼320 μm^2^ and 4× averaging was performed to improve the signal-to-noise ratio. Images were denoised using the Despeckle and Remove Outlier (3 pixel radius, threshold = 20) functions in Fiji (28).

### Drugs

Stock solutions of all drugs were diluted in standard Ames’ medium and constantly gassed with 95% O2 / 5% CO2 immediately prior to recording. All stocks were diluted in either Milli-Q water or DMSO depending on their solubility; stock solutions were at least 1000x of the final concentration in the bath. Below is a list of the drugs used in this experiment and delivered via a gravity fed perfusion system at 2–5 ml/min: Dopamine hydrochloride (Sigma-Aldrich; Saint Louis, MO), forskolin (Tocris; Bristol, UK), SKF383938 (Tocris; Bristol, UK), SCH39166 (Tocris; Bristol, UK), morphine sulfate salt pentahydrate (Sigma-Aldrich; Saint Louis, MO), [D-Ala2, NMe-Phe4, Gly-ol5]-enkephalin (DAMGO; Tocris; Bristol, UK), Naloxone (Tocris; Bristol, UK), and SR95531 hydrobromide (Tocris; Bristol, UK). Notably, ascorbic acid (Sigma-Aldrich; Saint Louis, MO) was added to dopamine stocks to prevent oxidation during the recording process. The final concentration in the recording solutions were 100 µM and 10 µM for dopamine hydrochloride and ascorbic acid, respectively. To inhibit PKA activity, 2.5 µM KT5720 (Tocris; Bristol, UK) was added to the CsGlc-based internal solution. GABA (100 µM, Sigma; St. Louis, MO) or glutamate (10 mM, Sigma; St. Louis, MO) was dissolved in standard bicarbonate-buffered extracellular Ames’ medium and was puff applied from small-tipped (∼1 µm diameter) glass pipettes positioned at ∼20–30 µm away from the ipRGCs with Picospritzer III (Parker, Cleveland, OH). GABA puffs at 8-10 psi for 1 second were aimed toward the ipRGCs, parallel to and in the direction of the flow of perfusion, while the cell was held at -70 mV.

### Data analysis

Data for GABA puff experiments were analyzed off-line using IgorPro software version 5.03 (Wavemetrics; Portland, OR). GABA puff-evoked currents were leak-subtracted and peak current amplitudes were measured for each trace. This data was then transferred into Microsoft Excel and imported to RStudio for data management, visualization, and analysis.

For the retinal whole mount experiment, spontaneous inhibitory postsynaptic currents (IPSCs) were detected using Mini analysis software (Synaptosoft; Fort Lee, NJ). While many ipRGCs had detectable inhibitory events, only cells that had a spontaneous IPSC frequency > 1 Hz in control conditions were included in the analysis. Data from the Mini Analysis software was then transferred into Microsoft Excel and imported to RStudio for data management, visualization, and final analysis.

Data are presented as mean ± standard error of the mean (SEM) as specified in figure captions. Prior to performing statistical tests, the data was assessed for normality of residuals as well as homogeneity of variance. Based on the data’s parameters, the data were then analyzed using a either t-test, Wilcoxon test or ANOVA depending on the dataset performed using RStudio (v2022.7.2.576) with Tukey’s post hoc adjustments for all pairwise comparisons (when applicable), and p < 0.05 was considered significant.

## RESULTS

### Inhibitory synaptic inputs to ipRGCs are primarily GABAergic

Previous data from our lab showed that MOR activation in ipRGCs reduces light-evoked firing (2). The use of dissociated retinal cultures and synaptic blocking cocktails for these previous experiments allowed for the examination of MOR activation on intrinsic excitability and light-responsiveness of ipRGCs. It is still unknown, however, whether retinal MOR activation might affect synaptic inputs to these ganglion cell photoreceptors (18). First, in whole mount retinas, we recorded both excitatory and inhibitory postsynaptic currents in the same ipRGCs in a consecutive manner, by alternately holding the membrane potential at -70 mV and 0 mV, respectively (29). Consistent with past studies (19), measurable fast excitatory (presumably glutamatergic) currents were detectable only in a very small subset of ipRGCs in the whole mount retina (n=3/58 cells, 5.2%). Additionally, even 10 mM glutamate puffed onto solitary ipRGCs resulted in no measurable current (data not shown). On the other hand, inhibitory postsynaptic currents (IPSCs) in EGFP-labelled ipRGCs in the whole mount retina held at 0 mV were regularly recorded (Fig. 1*A*). Since the reversal potential for chloride under these recording conditions was approximately -70 mV, IPSCs manifested as outward currents (Fig. 1*C*). Notably, the majority of these IPSCs were blocked by the application of the GABA_A_ receptor-selective antagonist Gabazine (SR-95531); which, consistent with previous studies (19), indicates that the inhibitory inputs to these ipRGCs are primarily GABAergic (Fig. 1). To measure GABAergic events more robustly in ipRGCs, we puffed 100 μM GABA for 1 second onto acutely dissociated ipRGCs using a CsGlc-based internal pipette solution with a higher chloride concentration (Fig. 1*B*). This allows us to consistently measure GABA puff-mediated currents at potentials that are closer to rest. Consistent with the idea that these puff evoked IPSCs are mediated by the GABA_A_ receptor, when we stepped the membrane to different potentials (-80 mV to 0 mV at 20 mV steps) and puffed GABA onto the cells, the resulting current reversed at around the calculated reversal potential for chloride (Fig. 1*D-E*). Moreover, this puff evoked current was blocked by the chloride channel blocker picrotoxin (100 μM), further suggesting that this puffing GABA onto solitary ipRGCs conducts chloride via the GABA_A_ receptor (Fig. 1*D-E*). Taken together, these data suggest that ipRGCs robustly express somatic GABA_A_ receptors which signal to synaptically modulate ipRGC excitability.

**Figure 1.**
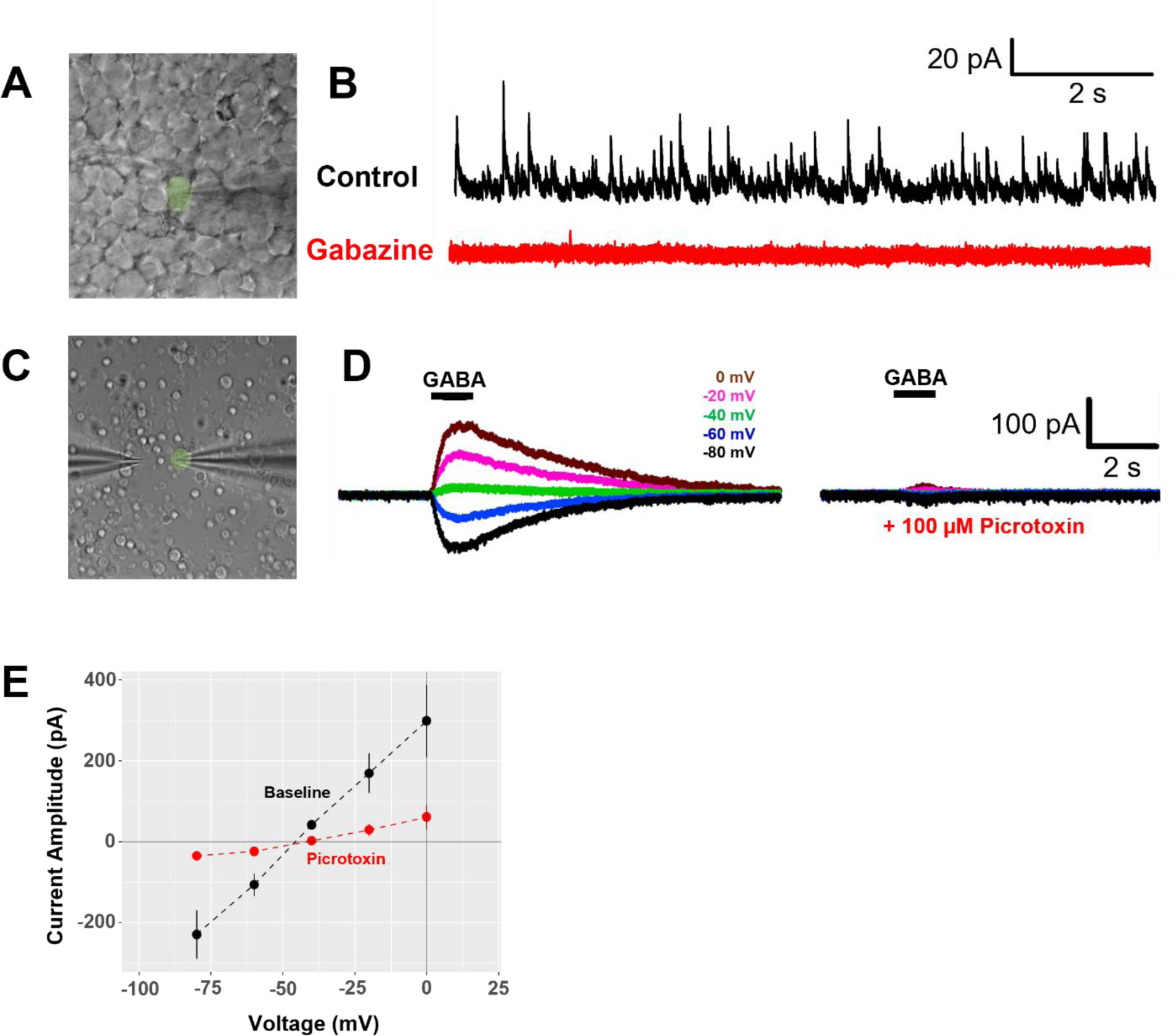
Inhibitory inputs to ipRGCs are primarily GABAergic. (A) *Opn4*::EGFP+ ipRGC in the whole mount retina with a patch pipette recording whole-cell synaptic currents. (B) Solitary *Opn4*::EGFP+ ipRGC in a dissociated retinal culture showing a patch pipette(right) recording whole-cell currents during application of 100 µM GABA by a puff pipette (left). (C) Spontaneous inhibitory events in EGFP+ ipRGCs are abolished by 30 µM SR-95531 (Gabazine) while holding the cell at + 0 mV. (D) Representative trace evoked by puffing 100 µM GABA onto solitary ipRGCs (left; D_1_) that were blocked by 100 µM picrotoxin (right; D_2_) at the following holding potentials: -80 mV (black), -60 mV (blue), -40 mV (green), -20 mV (pink), 0 mV (brown). (E) I-V curve cumulative data from GABA-puff evoked currents at different holding potentials (-80 mV to 0 mV in 20 mV steps) in control (n=6) and after the application of 100 µM picrotoxin (n=4).

### MOR activation does not affect GABA conductance in solitary ipRGCs

Given the variable results in the whole mount retina, we decided to assess how neuromodulators might alter GABAergic inputs to ipRGCs. We did this by puffing GABA directly onto acutely dissociated ipRGCs, which resulted in inward currents when the cell was held at -70 mV (Fig. 2). Indeed, there is significant evidence for opioidergic modulation of GABAergic signaling in the brain (21, 22). Therefore, we tested whether MOR activation altered somatic GABA currents (I_GABA_) in solitary ipRGCs. Accordingly, we compared the amplitudes of the initial puff-evoked I_GABA_ (control), to those recorded when the cells had been exposed to the MOR agonists, morphine or DAMGO (Fig. 2, *A-D*). First, we examined the amplitude of the puff-evoked I_GABA_ in control, after 5 min in 10 μM morphine and then following washout. Analysis of these currents using a one-way ANOVA revealed no significant changes in current amplitude across treatments (Fig. 2*C*, One-way ANOVA, *P =* 0.8777). A similar experiment was then performed by bath applying 1 μM of the MOR-selective agonist DAMGO, followed by perfusing 1 μM naloxone, a non-selective opioid receptor antagonist while 100 μM GABA was puffed onto solitary ipRGCs (Fig. 2*B*). Despite the use of a more selective MOR agonist (DAMGO), results of these experiments were similar to those observed following morphine application; with MOR activation yielding no significant change in the GABA-puff evoked current (Fig. 2*D*, One-way ANOVA, *P =* 0.7766). These indicate that MOR activation in ipRGCs does not affect the somatic I_GABA_ in solitary ipRGCs.

**Figure 2.**
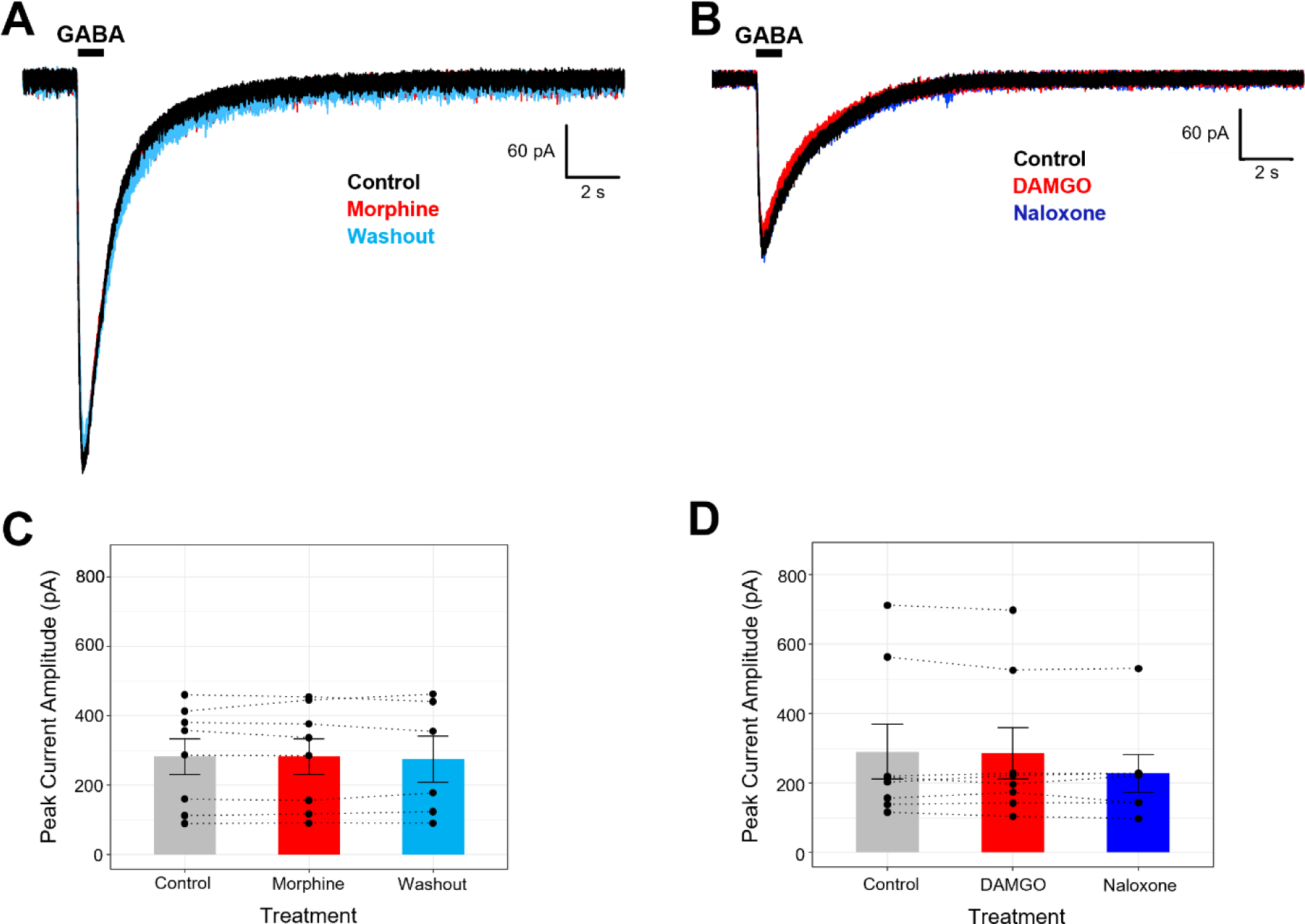
MOR activation does not affect puff-evoked GABAergic currents in solitary ipRGCs. (A) Representative trace evoked by puffing 100 µM GABA for 1 second onto solitary ipRGCs that were unaffected by application of 10 µM morphine. (B) Representative trace evoked by puffing 100 µM GABA onto solitary ipRGCs that were unaffected by application of 1 µM DAMGO or recovery with 1 µM Naloxone. (C) Cumulative data for solitary ipRGCs that were non-responsive in terms of the effect of morphine on the puff-evoked GABAergic IPSC (n=8). (D) Cumulative data for solitary ipRGCs that were non-responsive in terms of the effect of DAMGO on the puff-evoked GABAergic IPSC (n=8). One-way ANOVA performed with a Tukey post-hoc comparison. Data presented as the mean ± SEM.

### Dopamine enhances the GABAergic conductance via dopamine D1R in solitary ipRGCs

Since activation of the MOR appears not to alter I_GABA_ in ipRGCs, we wanted to assess whether dopamine, an important neuromodulator for light adaptation, alters the magnitude of GABA currents in ipRGCs. Not only have DAC been shown to corelease dopamine and GABA (30), but dopamine can also act to enhance GABAergic currents in retinal amacrine cells (31). Accordingly, we recorded puff-evoked GABA currents in ipRGCs in the presence of dopamine and/or the D1R-selective antagonist SCH39166. These experiments revealed that half of the cells recorded (6/12 cells) exhibited dopamine-mediated increases in GABA current amplitude, which was reversible by 10 µM SCH39166, while the other half were unaffected by dopamine application (Fig. 3, *A-D*, Dopamine non-responsive: control vs. dopamine: *P =* 0.9639; control vs. SCH39166: *P =* 0.9833; dopamine vs. SCH39166: *P =* 0.9962; Dopamine responsive: control vs. dopamine: *P <* 0.0001; control vs. SCH39166: *P =* 0.9711; dopamine vs. SCH39166: *P <* 0.0001, 2-way repeated measures ANOVA with Tukey posthoc tests). When comparing the dopamine-mediated increase in GABA current, dopamine-sensitive neurons exhibited a 28% mean increase in peak current amplitude compared to a 0.65% increase in dopamine non-responsive cells (Fig. 3*E*, *P* = 0.002, Wilcoxon rank-sum test).

**Figure 3.**
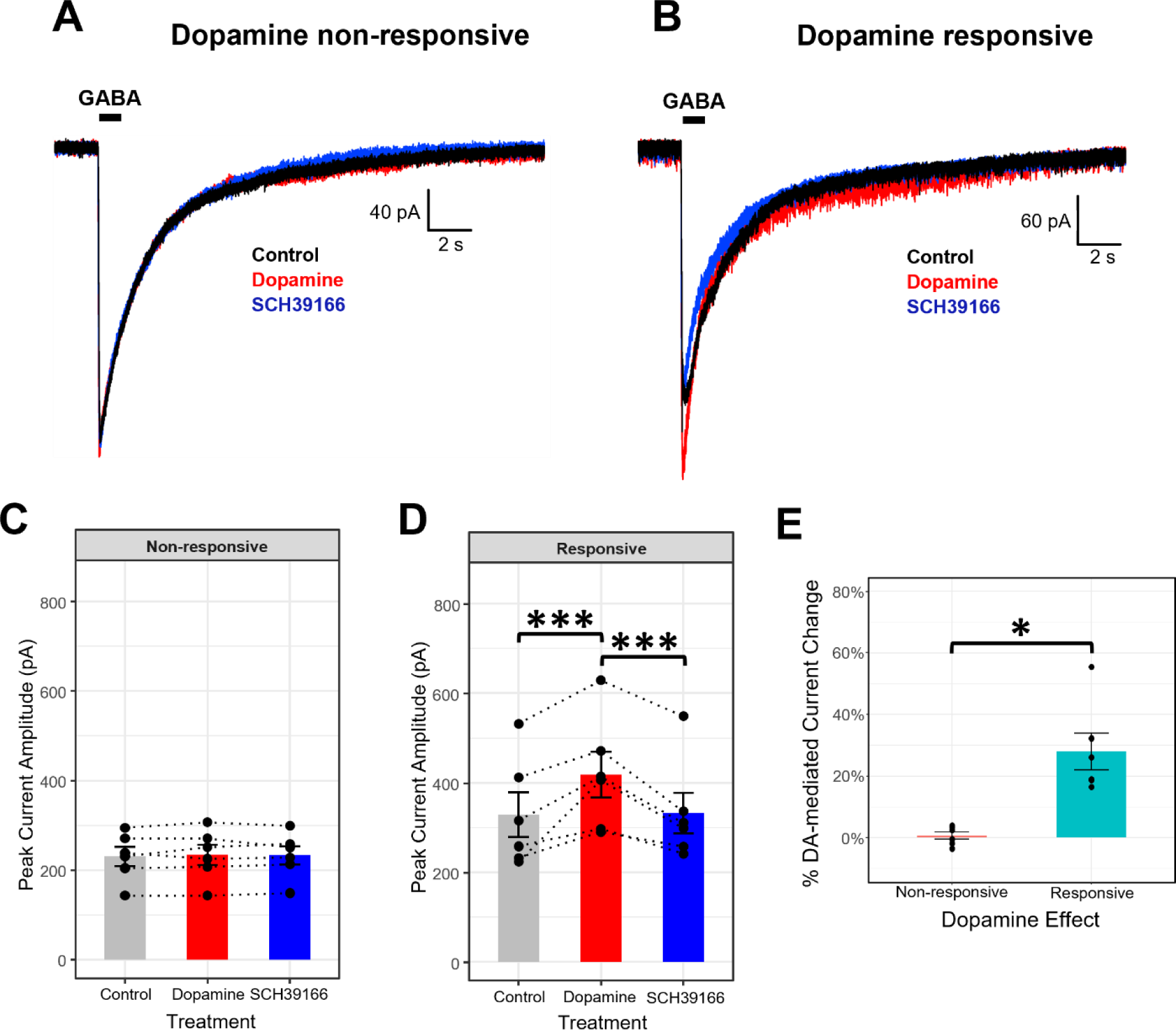
Differential responsiveness of puff-evoked GABAergic currents to dopamine in solitary ipRGCs. (A) Representative trace evoked by puffing 100 µM GABA for 1 second onto solitary ipRGCs that were unaffected by dopamine applied to the bath solution. (B) Representative trace evoked by puffing 100 µM GABA for 1 second onto solitary ipRGCs wherein dopamine enhanced the GABA-evoked current. (C) Cumulative data for 6 solitary ipRGCs that were non-responsive in terms of the effect of dopamine on the GABAergic IPSC. (D) Cumulative data showing 6 solitary ipRGCs wherein 100 µM dopamine increased the GABAergic IPSC and this effect was reversed by 10 µM of the D1R antagonist SCH39166. Two-way ANOVA performed with a Tukey post-hoc comparison. (E) Dopamine-responsive cells exhibit significant increases in the GABA-puff IPSC following dopamine exposure. Unpaired two-sample Wilcoxon test. Data presented as the mean ± SEM. (*: p<0.01, ***: p<0.0001)

Since dopamine can be washed-out in our dissociated retina preparation (Supp Fig. 1*B*), we were unsure whether the SCH39166 recovery of the GABA-evoked current was due to D1R antagonism or dopamine washout. To confirm that the facilitatory effect on the GABA current was mediated by D1R, we performed a similar experiment using the D1R-selective agonist SKF383938 (which we were unable to washout; Supp Fig. 1*C*). This experiment yielded similar results to the previous dopamine experiment, showing D1R-mediated facilitation of GABA currents in half of the recorded cells (n=7/14 cells). Here, the increase in GABAergic currents in SKF383938 responsive cells was reversible by SCH39166, while D1R agonist/antagonist did not affect SKF383938 non-responsive cells (Fig. 4, *A-D*, SKF383938 non-responsive: control vs. SKF383938: *P =* 0.4231; control vs. SCH39166: *P =* 0.7055; SKF383938 vs. SCH39166: *P =* 0.8846; SKF383938 responsive: control vs. SKF383938: *P <* 0.0001; control vs. SCH39166: *P =* 0.6611; SKF383938 vs. SCH39166: *P <* 0.0001, 2-way repeated measures ANOVA with Tukey posthoc tests). When comparing the D1R-dependent facilitation of SKF383938 sensitive vs. insensitive solitary ipRGCs, SKF383938 increased the I_GABA_ by 24.6% in responsive cells, with little change (-3.2%) in cells unresponsive to the D1R-selective agonist (Fig. 4*E*, *P* < 0.0001, Welsh’s two sample t-test).

**Figure 4.**
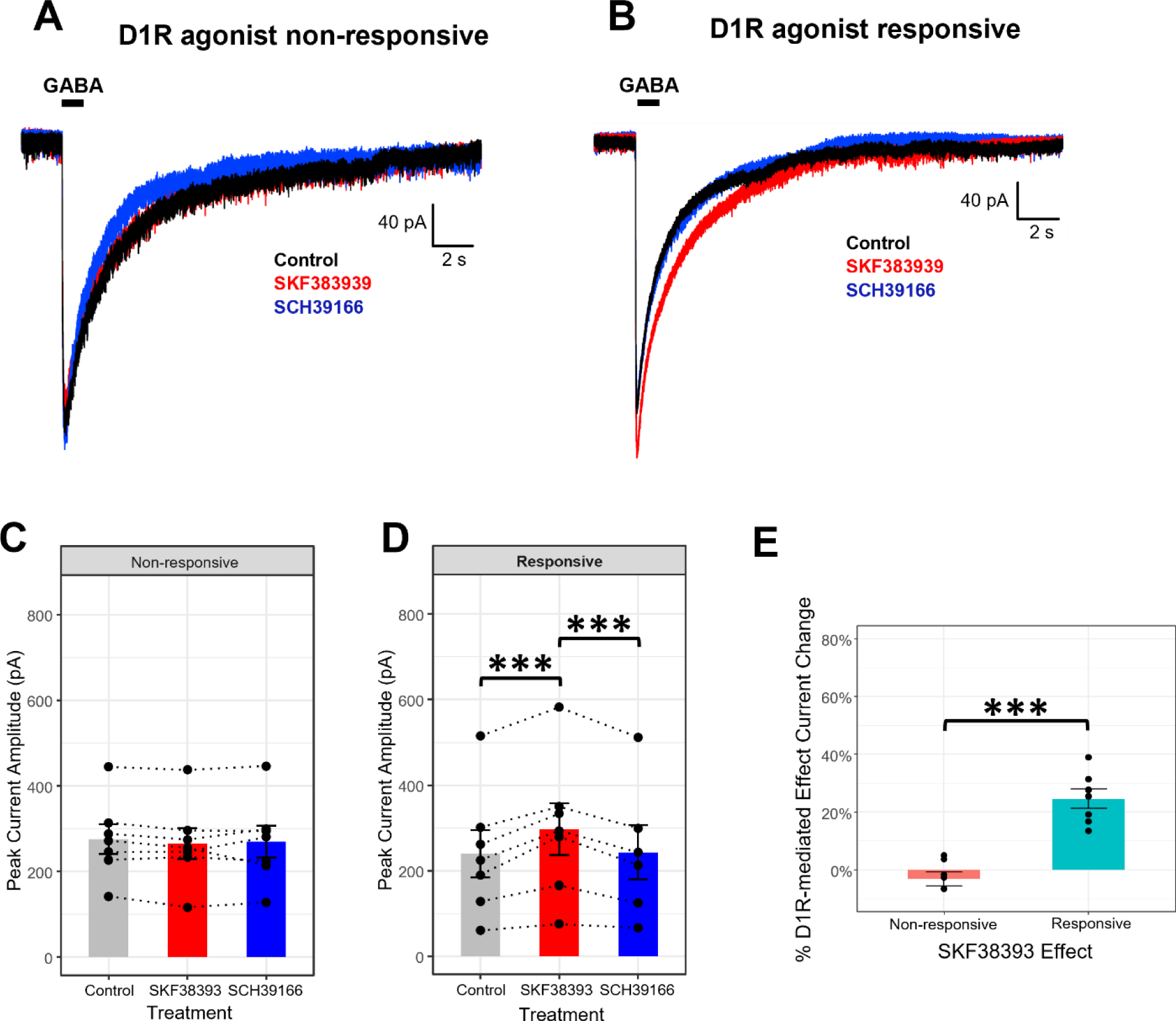
Differential responsiveness of puff-evoked GABAergic currents to D1R selective agonist SKF383939 in solitary ipRGCs. (A) Representative trace evoked by puffing 100 µM GABA onto solitary ipRGCs that were unaffected by 50 µM SKF383939 applied to the bath solution. (B) Representative trace evoked by puffing 100 µM GABA onto solitary ipRGCs wherein by 50 µM SKF383939 had a faciliatory effect on the GABAergic current. (C) Cumulative data for 7 solitary ipRGCs that were non-responsive in terms of the effect of SKF383939 on the GABAergic IPSC. (D) Cumulative data showing 7 solitary ipRGCs wherein 50 µM SKF383939 increased the GABAergic IPSC and this effect was reversed by 10 µM of the D1R antagonist SCH39166. Two-way ANOVA performed with a Tukey post-hoc comparison. (E) SKF383939 -responsive cells exhibit significant increases in the GABA-puff IPSC following D1R agonist exposure. Unpaired two-sample t-test. Data presented as the mean ± SEM. (***: p<0.0001)

Previous single-cell RT-PCR and immunohistochemistry experiments in rat showed that the majority of ipRGCs express the D1R (7). Here, we evaluated the prevalence of D1R expression in EGFP-expressing ipRGCs from *Opn4::*EGFP mice (n=5). In each mouse retina, we identified EGFP+ ipRGCs that were immunopositive for D1R along with ipRGCs that were not immunoreactive for the D1R (Fig. 5, *A-F*). However, across all the mice, we found that only 14 of the 121 EGFP+ ipRGCs showed positive D1R immunolabeling (11.6%) (Table 1). This finding shows that while there are both D1R-expressing and non-D1R-expressing ipRGCs, the prevalence of D1R expression in mouse ipRGCs is potentially reduced compared to expression within the rat retina (7).

**Figure 5.**
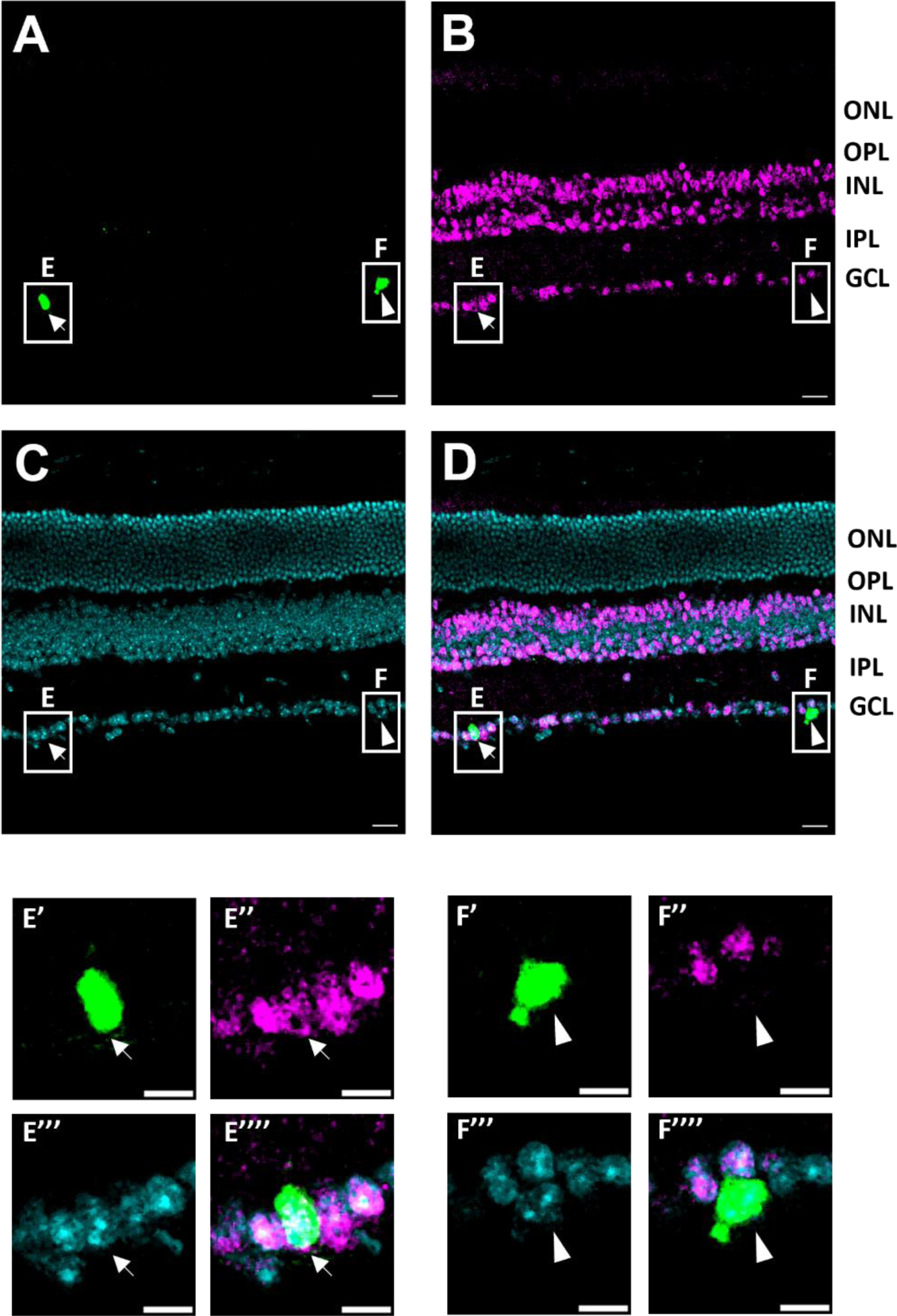
The dopamine receptor 1 (D1R) is expressed in a subset of *Opn::*EGFP+ ipRGCs. The D1R was immunolabeled (magenta, B) in *Opn::*EGFP+ (green, A) mice (n=5). Representative single optical section of a cryosectioned retina wherein only some EGFP-expressing ipRGCs are immunopositive for D1R (arrow, E’-E’’’) while others are negative (arrowhead, F’-F’’’’). Nuclei were stained with DAPI (cyan, C). Scale bars in large images (A-D) = 20 μm; scale bars in small images (E’-E’’’, F’-F’’’’) = 10 μm. ONL=outer nuclear layer; OPL= outer plexiform layer; INL: inner nuclear layer; IPL: inner plexiform layer; GCL: ganglion cell layer.

**Table 1.** Summary data for immunohistochemical detection of D1R expression in EGFP+ ipRGCs.

| Mouse ID | Sex | # D1R+ ipRGCs | Total # ipRGCs | % ipRGCs expressing D1R |
| --- | --- | --- | --- | --- |
| #64 | male | 1 | 25 | 4% |
| #103 | male | 2 | 16 | 12.5% |
| #104 | male | 3 | 7 | 43% |
| #72 | female | 3 | 33 | 9% |
| #78 | female | 5 | 40 | 12.5% |
| totals |  | 14 | 121 (11.6%) | 16.2% (average) |

Next, we tested how dopamine affects inhibitory inputs to ipRGCs *in situ*, in the whole mount retina. Thus, we recorded spontaneous inhibitory events from EGFP+ ipRGCs in the retinal whole mount while holding the cells at +0 mV to minimize cation flow through mixed cation channels. In this experiment, dopamine had no effect on the IPSC amplitude, whereas SCH39166 significantly decreased the amplitude of inhibitory events (Fig. 6*B*, control vs. dopamine: *P =* 0.1435; control vs. SCH39166: *P =* 0.0001; dopamine vs. SCH39166: *P =* 0.0017, One-way ANOVA with Tukey posthoc tests). Additionally, SCH39166, but not dopamine decreased the frequency of spontaneous inhibitory currents compared to control (Fig. 6*B*, control vs. dopamine: *P =* 0.9889; control vs. SCH39166: *P =* 0.0479; dopamine vs. SCH39166: *P =* 0.0379, One-way ANOVA with Tukey posthoc tests). These results are consistent with the idea that dopamine cannot be washed out of the light adapted retina (32). In the light, dopamine receptors are activated by endogenous dopamine released from DACs (13), such that exogenous bath-applied dopamine may not have an effect. In this way, only the application of D1R antagonist would reverse any synaptic effects that dopamine might have on neurons in the light-adapted retina. Taken together, these data suggest that a subset of ipRGCs express the D1R, which can enhance somatic GABA inputs to a specific population of ipRGCs.

**Figure 6.**
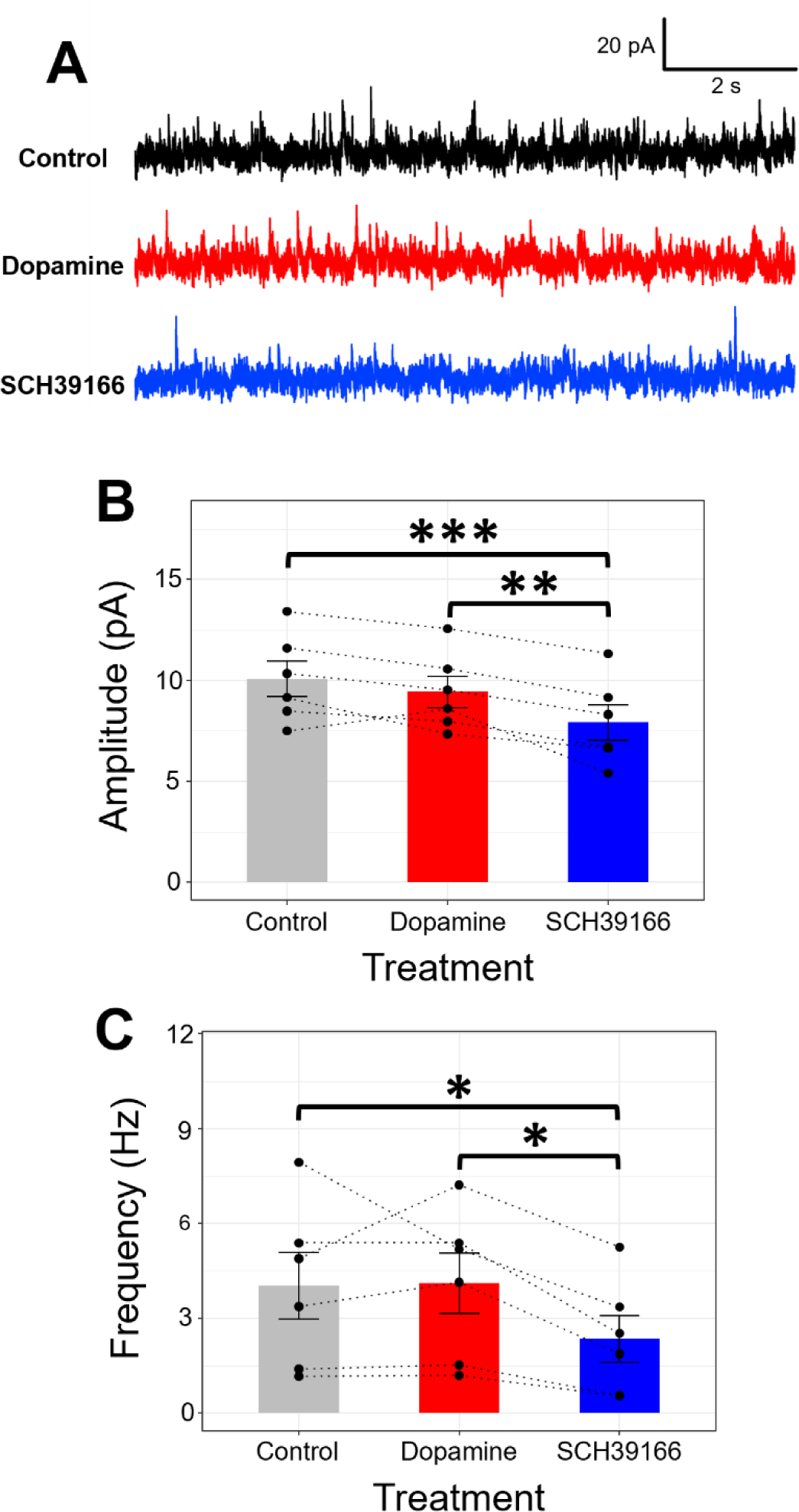
Dopamine D1 receptor (D1R) antagonism decreases inhibitory inputs to ipRGCs. (A) Representative traces showing decreased inhibitory events following application of 10 μM D1R antagonist SCH39166. (B) D1R antagonist SCH39166 decreased the amplitude of inhibitory inputs to ipRGCs. (C) D1R antagonist SCH39166 significantly decreased the frequency of inhibitory inputs to ipRGCs. One-way ANOVA performed with a Tukey post-hoc comparison. (*: p<0.05, **p<0.001, ***p<0.001; n=6 cells). Data presented as the mean ± SEM.

### Dopaminergic facilitation of somatic GABA currents occurs via a cAMP-PKA dependent mechanism

Given the importance of dopamine in retinal light adaption and motivated behaviors, much research has focused on D1R signaling (9). Canonically, the D1R is a Gαs-coupled receptor which alters intracellular functions primarily by activating adenylate cyclase following agonist binding and G protein dissociation (33). Previous studies have shown a D1R-mediated facilitation of GABAergic currents in retinal neurons via a PKA-dependent mechanism (31). To test whether this pathway is responsible for the D1R-mediated GABAergic facilitation in ipRGCs, we first attempted to recapitulate the dopamine-induced increase in I_GABA_ by directly stimulating the primary downstream effector of the D1R: adenylyl cyclase (AC). First, we had to determine the concentration of forskolin (a potent AC activator) that was appropriate for the proposed experiment. We determined that 5 μM forskolin was insufficient to enhance I_GABA_, but 15.8 μM and 50 μM forskolin sufficiently increased GABA conductance (Fig. 7 & Fig. S2).

**Figure 7.**
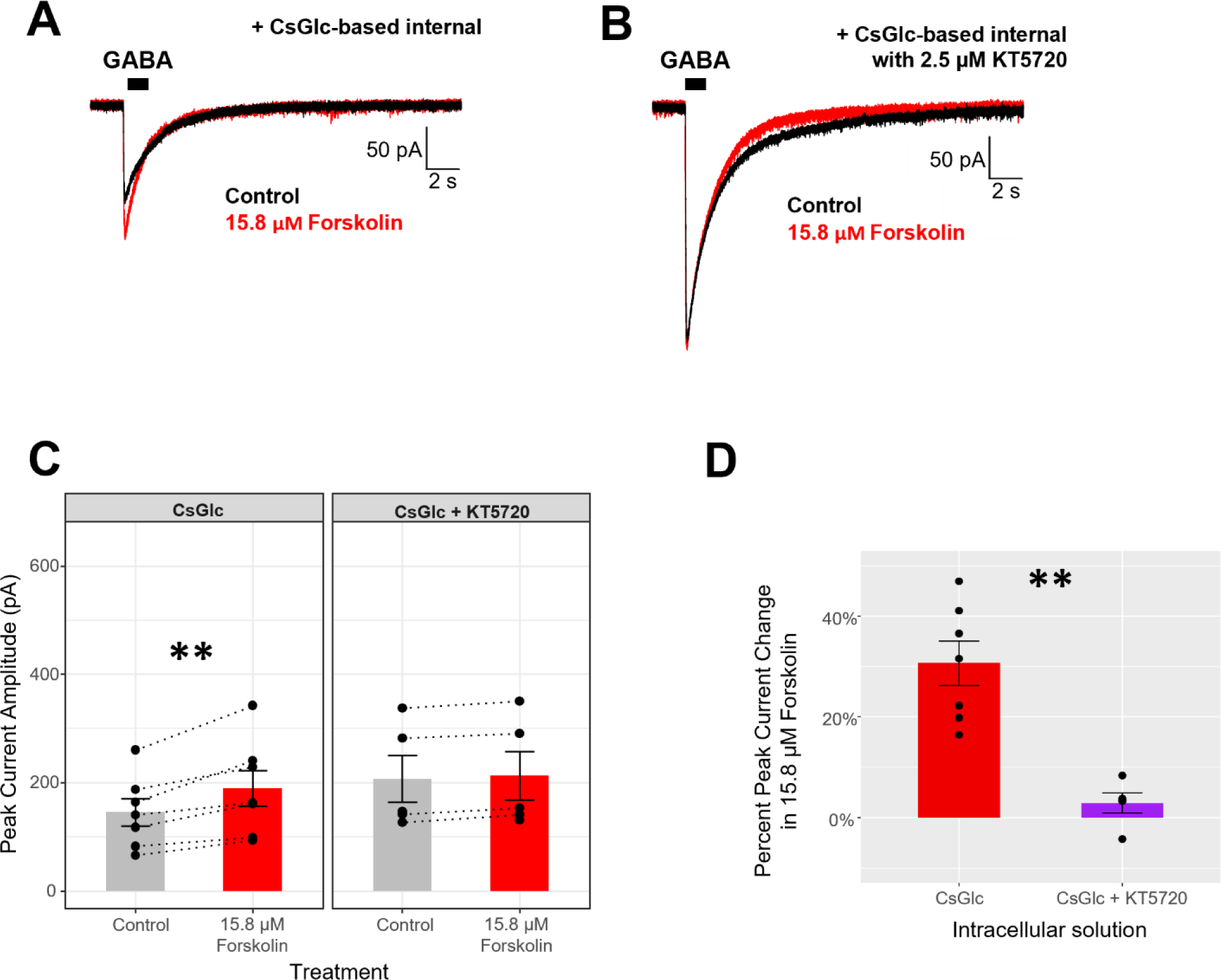
Direct activation of adenylate cyclase enhances the GABAergic IPSC in solitary ipRGCs via a PKA-dependent mechanism. (A) Representative trace showing that 15.8 µM forskolin enhanced the GABA puff-evoked IPSC in solitary ipRGCs. (B) Representative trace showing that 2.5 µM KT5720 blocked the 15.8 µM forskolin-induced increase in the GABA puff-evoked IPSC. (C) Cumulative data showing that forskolin-mediated increase in the GABAergic IPSC is prevented by intracellular application of the PKA inhibitor KT5720 (CsGlc: n=7; CsGlc+KT5720: n=5 Two-way ANOVA with Tukey posthoc tests on pairwise comparisons). (D) KT5720 abolishes the 15.8 µM forskolin-induced facilitation of GABA conductance in ipRGCs. Unpaired two-sample t-test. Data presented as the mean ± SEM. (**=p<0.001)

We attempted to block forskolin’s faciliatory effect on GABA_A_ receptor conductance by inhibiting Protein Kinase A (PKA), a canonical downstream target of AC/cAMP. Unfortunately, we were unable to wash forskolin out of the dissociated retinal culture. Thus, to test PKA’s involvement in D1R-mediated GABAergic facilitation in ipRGCs, we applied forskolin to solitary ipRGCs while using a pipette solution containing 2.5 μM of the PKA inhibitor KT5820. Interestingly, the PKA inhibitor KT5720 applied to the standard CsGlc-based internal recording solution abolished the 15.8 μM (but not the 50 μM) forskolin-induced increase in the GABA puff-induced IPSC (Fig. 7*C* & Fig. S2, CsGlc: control – 15.8 μM forskolin: *P =* 0.0002; CsGlc+KT5720: control – 15.8 μM forskolin: *P =* 0.5132, Two-way repeated measures ANOVA). In line with this, the 30.6% increase in I_GABA_ induced by 15.8 μM forskolin was significantly reduced to just 2.9% in the presence of KT5720 (Fig. 7*D*, CsGlc – CsGlc+KT5720: *P* = 0.00037, Welsh’s two sample t-test). Additionally, when comparing the forskolin-mediated increase in the GABAergic peak current, both 5 μM forskolin and 15.8 μM with KT5820 in the internal solution did not significantly increase the GABAergic IPSC, while 15.8 μM and 50 μM forskolin increased the GABA current by 30% and 28.7%, respectively (Fig. S2*F,* 5 μM forskolin – 15.8 μM forskolin: *P <* 0.0001, 5 μM forskolin – 50 μM forskolin: *P <* 0.0001, 50 μM forskolin – 15.8 μM forskolin: *P =* 0.9670, 5 μM forskolin – 15.8 μM forskolin + KT5720: *P =* 0.6132, 15.8 μM forskolin – 15.8 μM forskolin + KT5720: *P <* 0.0001, 50 μM forskolin – 15.8 μM forskolin + KT5720: *P =* 0.0001, One-way ANOVA with Tukey posthoc corrections). These data point to a cAMP-PKA dependent facilitation of GABA_A_ receptor-mediated conductance in ipRGCs.

## DISCUSSION

### Putative mechanism of D1R-mediated facilitation of GABA_A_ receptor conductance

Based on the electrophysiological and immunohistochemical data presented in this paper, D1R activation enhances the GABA_A_ receptor-mediated currents in a subset of ipRGCs. A similar effect has been shown in amacrine cells in which D1R activation drastically increases the I_GABA_ via a cAMP/PKA-dependent pathway (31). While the D1R-mediated increase of the I_GABA_ through the GABA_A_ receptor is not quite as robust in ipRGCs as in amacrine cells (31), the data presented here provides strong evidence that dopamine-mediated postsynaptic facilitation of I_GABA_ in ipRGCs occurs through a cAMP-PKA dependent mechanism. It is well established that Gαs-coupled receptors such as the D1R undergo a conformational change upon agonist binding that leads to GTP/GDP exchange and dissociation of G protein subunits (33). The α subunit then is free to activate AC, whose catalytic activity increases intracellular cAMP levels (Fig. 8). Indeed, we show that increased AC activity (with forskolin) mimics D1R agonism in terms of GABAergic current facilitation. Notably, the forskolin effect occurred in all recorded EGFP+ ipRGCs rather than a subset (as with dopamine/SKF383938), presumably because AC is expressed in all ipRGCs (34). AC activation and the subsequent increases in intracellular cAMP levels promote cAMP binding to the regulatory subunits of cAMP; allowing for PKA to carry out catalytic activity on downstream effectors (Fig. 8). Importantly, PKA has been shown to have differential effects on GABAergic currents. In fact, the effect that PKA has on GABA_A_ receptor-mediated conductance is largely dependent on the β subunit composition of the GABA_A_ receptor itself. For instance, GABA_A_ receptors containing both β_1_ and β_3_ subunits have been shown to decrease their conductance following phosphorylation by PKA (35). On the other hand, GABA_A_ receptors whose β subunits are both of the β_3_ variety increase GABA_A_ receptor-mediated currents through PKA-dependent phosphorylation in retinal neurons and brain cells alike (31, 35, 36). The fact that PKA inhibition effectively blocked the forskolin-mediated enhancement of GABA currents in ipRGCs, suggests the GABA_A_ receptor expressed by ipRGC are β_3_ subunit-containing pentamers (31, 32, 37, 38). While it is possible that D1R signaling through cAMP/PKA causes recruitment of additional GABA_A_ receptors to the membrane to pass additional current, there is little empirical evidence to back this idea in the context of GABA_A_ receptor signaling in ipRGCs. Therefore, the D1R-mediated enhancement of GABA currents (through cAMP/PKA) favors the idea that the phosphorylation of the GABA_A_ receptor increases its conductance for chloride.

**Figure 8.**
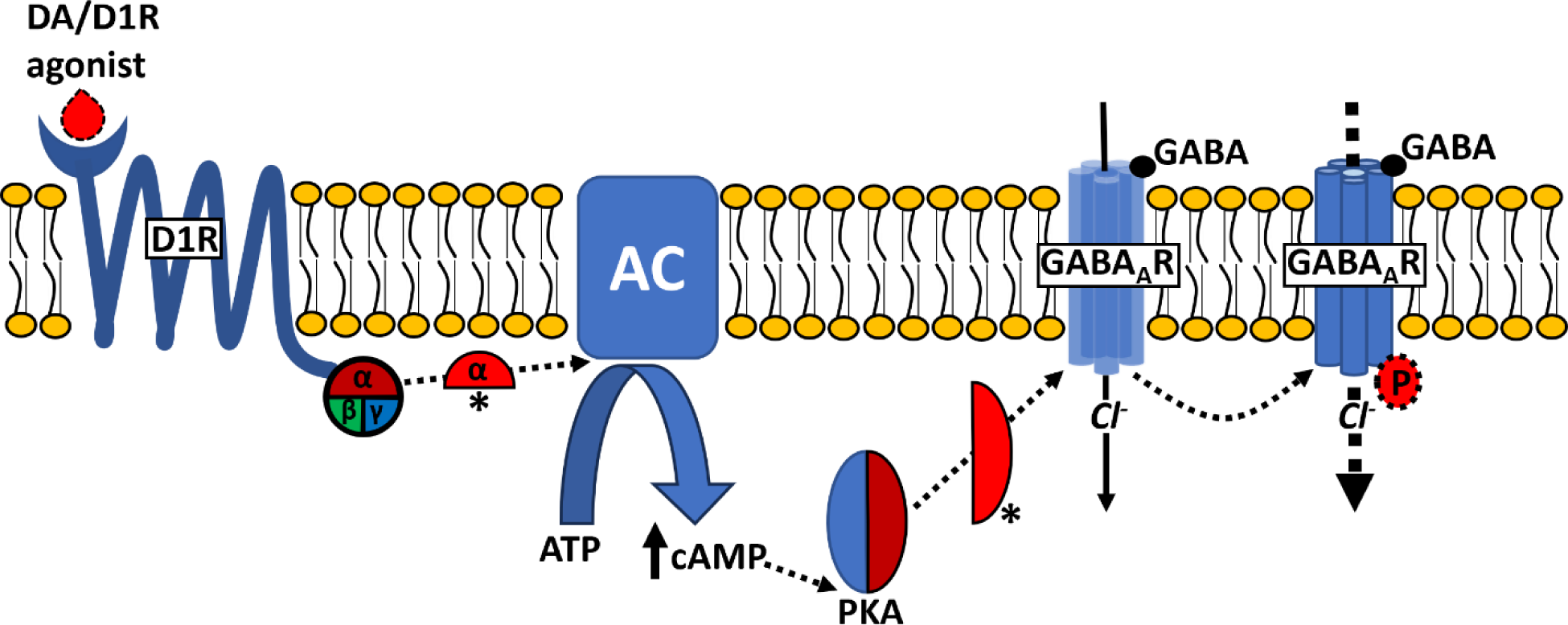
Putative pathway underlying D1R-mediated facilitation of GABA_A_ receptor currents in ipRGCs. D1R activation causes a conformational change that leads to dissociation of the connected G protein, leading to activation of adenylate cyclase (AC) via the α subunit. AC activation subsequently increases intracellular levels of cAMP which can bind to the regulatory subunits of Protein Kinase A (PKA). cAMP binding to PKA activates the catalytic subunits of PKA which can phosphorylate the GABA_A_ receptor (GABA_A_R), allowing for increased chloride (Cl^-^) conductance upon agonist exposure.

### The role of dopamine/D1R signaling in tuning ipRGC excitability

GABA puff experiments show that dopamine can significantly increase the I_GABA_ in specific ipRGCs, however, the same was not evident when recording spontaneous IPSC from the whole mount retina. While dopamine can be effectively washed out of an acutely dissociated culture (Fig. S1*B*), the same is not true for a retinal whole mount preparation. Here, application of 100 µM dopamine showed no effect on GABAergic IPSCs amplitude/frequency compared to control (Fig. 6). This is likely because dopamine levels in the light adapted retina are already saturated. Indeed, light adapted retinas contained sufficient concentrations of dopamine to activate the D1R (39, 40); thus, the addition of exogenous dopamine would have no effect on the amplitude and frequency of spontaneous IPSCs. Additionally, previous studies have shown that dopamine cannot be washed out of the whole mount retina (32). This might suggest that retinal dopamine receptors are already maximally activated prior to the application of D1R antagonist SCH39166. Within this context, it follows that D1R antagonism would decrease spontaneous IPSC amplitude, effectively blocking the facilitatory effect of endogenous light induced D1R activation on facilitation of GABA_A_ receptor-dependent currents.

Interestingly, bath application of SCH39166 also decreased the frequency of spontaneous IPSCs in the whole mount preparation. The interpretation of these data is less straightforward, however, dopamine also affects cells presynaptic to DACs as well as DAC coupling with other retinal neurons (4, 41). Indeed, D1R activation is associated with decreased coupling (via gap junctions) to other amacrine cells (41). Thus, D1R antagonism could increase amacrine cell coupling, which has been shown to lower circuit activity (42) and reduces resistance of individual amacrine cells within this coupled system. The reduced input resistance of the coupled GABAergic amacrine cells would result in smaller depolarization in response to excitatory inputs and in turn reduced GABA release onto postsynaptic targets such as ipRGCs. This might explain the decrease in spontaneous presynaptic GABA release detected in ipRGCs in the presence of the D1R antagonist SCH39166.

D1R receptor signaling within ipRGCs appears to enhance the postsynaptic GABA effect in a subset of ipRGCs. In the acutely dissociated retinal cultures, half the cells were responsive to dopamine/D1R agonist SKF383938 (Figure 2 & 3); but only 16% of EGFP+ ipRGCs appeared to express the D1R (Table 1). Previous work examining light-responsiveness of ipRGCs did not report a divergence between cells in terms of the responsiveness to dopamine and also detected a far greater proportion of D1R+ ipRGCs (7). This discrepancy between past findings and our data could have resulted from variations in tissue preparation as well as species differences (rat vs. mouse, respectively). While antibodies directed against receptor proteins, particularly those against G protein-coupled receptors, are notoriously unreliable (43), previous studies using the same antibody detected the D1R in a majority of M1 ipRGCs (7). Although we targeted M1 ipRGCs (based on size and fluorescence, see Materials and Methods) for immunohistochemistry and electrophysiology, the *Opn4*::EGFP line has been reported to label M1-M3 subtypes of ipRGCs (23). It is possible that D1R abundance could be higher in the M1 subtype of ipRGCs compared to M2-M3 subtypes, which gave us a lower number of D1R expressing/dopamine responsive cells compared to previous investigations. Another possible explanation for the discrepancy between imaging and physiology might be explained by the potential activation of the D5-subclass of dopamine receptors (D5R) in ipRGCs. Indeed, D5R receptors are characterized as D1-like receptors that can activated by the D1R agonist SKF383938 and blocked by the D1R antagonist SCH39166 (44–47).

Along these same lines, it would be interesting to determine whether the projection regions that dopamine-responsive ipRGCs differ from ipRGCs that lack functional D1-like receptors. Indeed, dopamine helps set the gain in the retina to allow for optimal functioning in bright light conditions (13). Yet, less is known about how retinal dopamine signaling affects downstream projection areas in the brain. Understanding the relationship between D1-like receptor expression in ipRGCs and their downstream projection area could further establish a role for retinal dopamine signaling within the context of both image forming and non-image forming visual functions. Indeed, drugs such as cocaine and amphetamines as well as Parkinsonian disorders can alter dopamine levels and disrupt the retina’s ability to adapt based on environmental illumination (4, 9). More specifically, alterations in retinal dopamine could modify ipRGC activity such that the light/dark signals transmitted to the brain are distorted. In this way, dysfunction in retinal dopamine signaling could explain some of the sleep/wake problems that are associated with dopamine-dependent stimulant use (48) as well as Parkinsonisms (49).

## Conclusions

The data presented in this paper show that some ipRGCs express D1-like receptors which can effectively modulate the strength of inhibitory synaptic inputs. In the context of dopamine-mediated light adaptation of the retina, it has been shown that dopamine enhances both excitatory and inhibitory synaptic currents alike (31, 32, 37, 38, 50). However, based on the data presented in this study, it appears that D1R-mediated signaling in ipRGCs plays a larger role in enhancing fast inhibitory synaptic inputs than affecting excitatory ones. When we consider that DACs co-release GABA and dopamine in the presence of bright light (30), it appears that dopamine functions to amplify the DAC-derived inhibitory inputs to ipRGCs. Interestingly, the mechanism by which D1-like receptor activation enhances GABA currents in ipRGCs, parallels and is synergistic to the proposed mechanism by which D1R activation attenuates melanopsin photocurrents and light-evoked ipRGC firing (7, 51). Indeed, D1R agonism is expected to lead to downstream (cAMP/PKA-dependent) phosphorylation of melanopsin to suppress light responsiveness of ipRGCs (51). Thus, D1-like receptor activation (via cAMP/PKA-mediated phosphorylation) increases inhibitory synaptic inputs, while concurrently suppressing melanopsin-based light responses to inhibit ipRGC firing in the presence of high retinal concentrations of dopamine. Although it is also possible that dopamine can signal through other receptors to affect ipRGC excitability, in light conditions when retinal dopamine concentrations are highest, D1-like receptor signaling is likely favored over D2-like receptor signaling pathways (40). It is, therefore, intriguing to consider that D1R signaling might function as a mechanism to prevent the overexcitation of ipRGCs in bright light conditions. To fully appreciate dopamine’s impact on ipRGC function, future studies could explore the role that dopamine has on other neuromodulatory inputs to ipRGCs.

## DATA AVAILABILITY

All data, including the code used for data analysis and visualization, is available at https://github.com/thebergular/D1R_GABA_ipRGCs.

## ACKNOWLEDGMENTS

We would like to thank Christopher Vaaga and Paul Witkovsky for their valuable input and review of the manuscript.

## AUTHOR CONTRIBUTIONS

Identify which authors participated in the research: N.B. and J.V. conceived and designed research; N.B. and C.-T.-B performed experiments, N.B. analyzed data, N.B. interpreted results of experiments, N.B. prepared figures, N.B. drafted manuscript, N.B. and J.V. edited and revised manuscript, approved final version of manuscript.

## FIGURE LEGENDS

**Figure S1.**
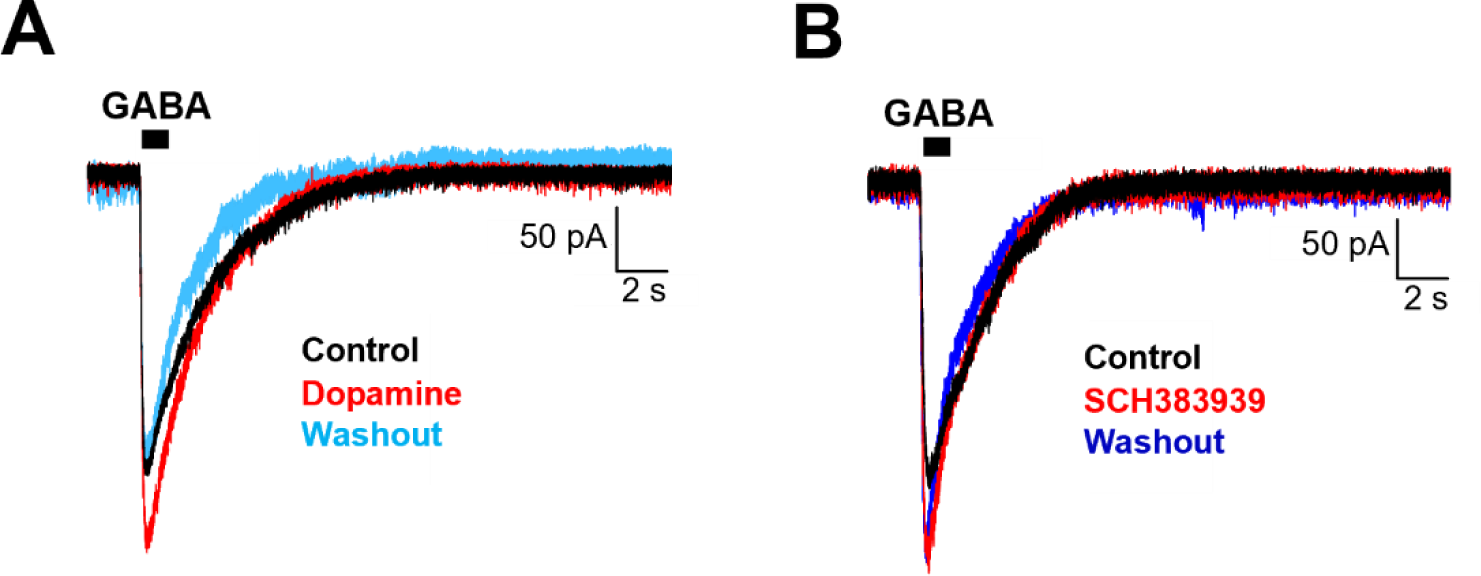
In acute retina cultures the dopamine effect, but not SCH383939 effect, on the GABAergic IPSC was reversible via washout. (A) Representative trace showing that 100 µM dopamine-mediated increase in the GABA puff-evoked IPSC was reversed following a 4-5 minute wash out. (B) Representative trace showing that 50 µM D1R agonist SCH383939 enhanced the GABA puff-evoked IPSC, which could not be reversed by drug washout after 10+ minutes.

**Figure S2.**
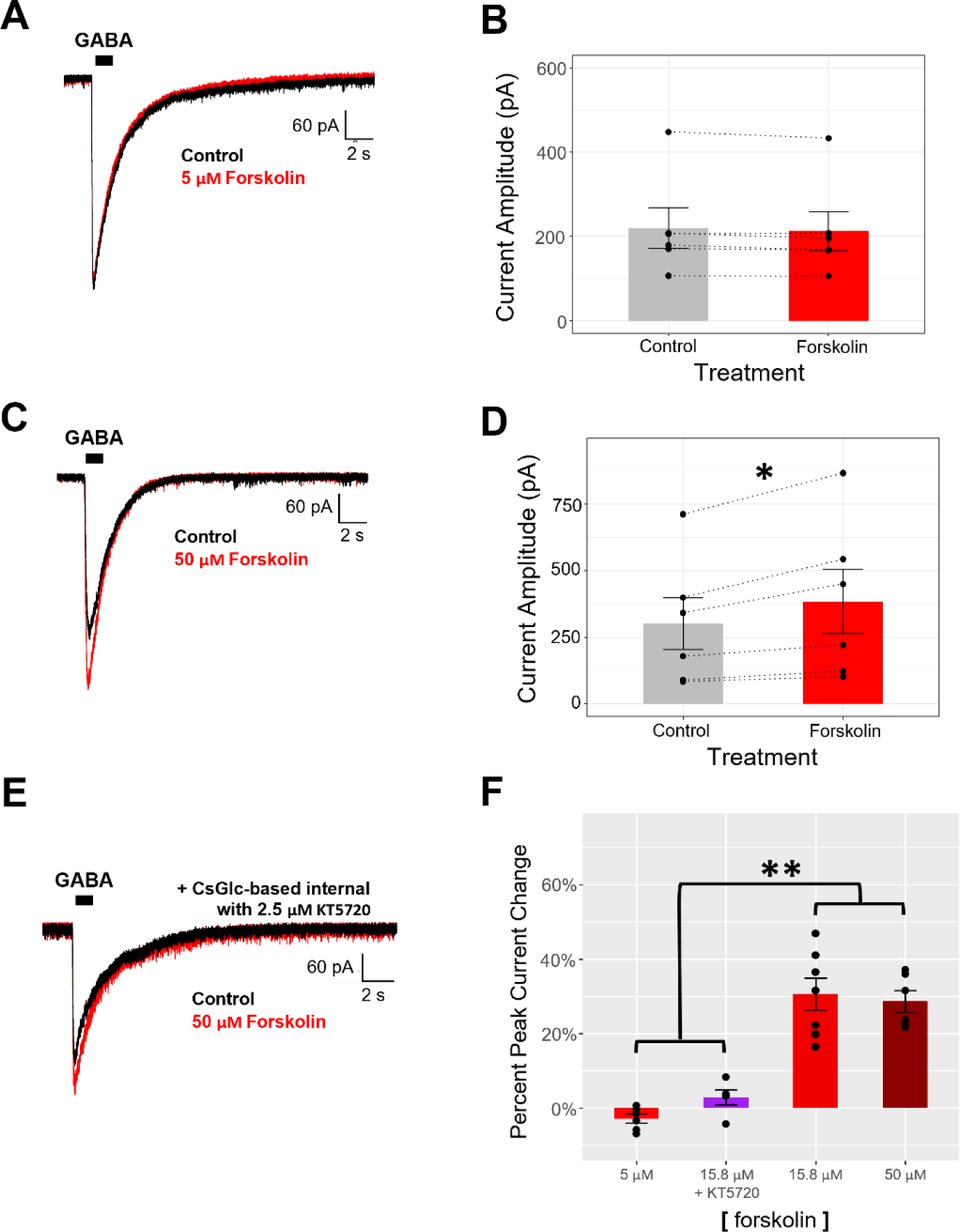
Optimization of forskolin concentration for enhancement of the GABAergic IPSCs in solitary ipRGCs. (A) Representative trace showing that 5 µM forskolin did not alter the GABA puff-evoked IPSC in solitary ipRGCs. (B) Summary statistics showing that the GABAergic IPSC was unaffected by the bath application of 5 µM forskolin (n=6, paired t-test). (C) Representative trace showing that 50 µM forskolin enhanced the GABA puff-evoked IPSC in solitary ipRGCs. (D) Summary statistics showing that the GABAergic IPSC is significantly enhanced by the bath application of 50 µM forskolin (n=6, paired t-test). (E) Representative trace showing that 50 µM forskolin enhanced the GABA puff-evoked IPSC in solitary ipRGCs, even in the presence of the PKA inhibitor KT5720 (2.5 µM in the standard CsGlc-based internal solution). (F) Comparisons of the peak current changes induced by different concentrations of forskolin. One-way ANOVA performed with Tukey posthoc test on all pairwise comparisons. Data presented as the mean ± SEM. (*=p<0.05, **=p<0.001)

